# Deep Learning for MYC binding site recognition

**DOI:** 10.1101/2022.03.03.482763

**Authors:** R. Fioresi, P. Demurtas, G. Perini

## Abstract

**Motivation:** The definition of the genome distribution of the Myc transcription factor is extremely important since it may help predict its transcriptional activity particularly in the context of cancer. Myc is among the most powerful oncogenes involved in the occurrence and development of more than 80% of different types of pediatric and adult cancers. Myc regulates thousands of genes which can be in part different, depending on the type of tissues and tumours. Myc distribution along the genome has been determined experimentally through chromatin immunoprecipitation This approach, although powerful, is very time consuming and cannot be routinely applied to tumours of individual patients. Thus, it becomes of paramount importance to develop in-silico tools that can effectively and rapidly predict its distribution on a given cell genome. New advanced computational tools (DeeperBind) can then be successfully employed to determine the function of Myc in a specific tumour, and may help to devise new directions and approaches to experiments first and personalized and more effective therapeutic treatments for a single patient later on.

**Results:** The use of DeeperBind with DeepRAM on Colab platform can effectively predict the binding sites for the MYC factor with an accuracy above 0.96 AUC, when trained with multiple cell lines. The analysis of the filters in DeeperBind trained models shows, besides the consensus sequence CACGTG classically associated to the MYC factor, also the other consensus sequences G/C box or TGGGA, respectively bound by the SP1 and MIZ-1 transcription factors, which are known to mediate the MYC repressive response. Overall, our findings suggest a stronger sinergy between the machine learning tools as DeeperBind and biological experiments, which may reduce the time consuming experiments by providing a direction to guide them.

## 1 Introduction

The myelocytomatosis, or MYC, gene belongs to a family of oncogenes coding for transcription factors involved in cell growth and in the activation and the expression of many pro-proliferative genes. In particular the MYC gene and its coded transcription factor lie at the crossroad of many signal transduction pathways and constitutes an early response downstream of many ligand-membrane receptor complexes, Armelin *et al*. (1984).

Given its important role, MYC expression is heavily regulated by several molecular mechanisms acting on many trascriptional regulatory motifs found within its proximal promoter region, Hurley *et al*. (2006). MYC over expression is very common in several diagnosed types of cancer in which it is expressed constitutively, Ahmadiyeh *et al*. (2010). A significant example is the Burkitt lymphoma in which MYC disregulations and mutations are well studied pathogenic causes, Bhatia et al. (1993), Magrath (1990). MYC encoded protein mechanism of action involves the formation of an heterodimer with the related transcription factor MAX, Blackwood and Eisenman (1991) (see Fig. 1 for a schematic picture of this mechanism). The heterodimer binds a 6-base DNA sequence (CACGTG) called E-box. If the binding occurs within a gene promoter and possibly nearby the transcription start site of the gene, it usually results in the activation of gene transcription. MYC/Max can also bind other DNA sequences which are slightly divergent from the canonical E-box such as CAtGTG, or CACGcG and have different affinity for the transcription factor, Eisenman (2001). Several studies have shown that the use of different genome DNA binding sequences may depend on the internal concentration of the Myc/Max complex, Sabo *et al*. (2014). In pathological condition in which the MYC gene is amplified and overexpressed, it has been found that Myc can also regulate transcription through a repressive mechanism, Herkert and Eilers (2010). However, in that case, it has been quite difficult to identify a specific DNA site recognized by the Myc/Max complex onto the genome. It rather seems that such repressive function is exerted by Myc by associating to chromatin through the interaction with other transcription factors. Several lines of evidence have shown that Myc can associate with either the Sp1 transcription factor or with the initiator Miz-1factor, Iraci *et al*. (2011). The interaction with SP1 or MIZ-1 allows Myc to recruit or regulate multiple chromatin modifiers and readers such as Histone Deacetylases, Sirtuins, Histone Demethylases and DNA Methyl Transferase; in other words, proteins involved in inducing heterochromatin formation to silence gene transcription, Lourenco *et al*. (2021). The possibility to predict binding of MYC to genomic DNA appears relatively easy, as it should be sufficient to bioinformatically identify the 6-base DNA sequences that can be recognized by the Myc/Max complex. However, sequence specificity alone cannot fully explain the MYC-MAX complex binding dynamic and occupancy across the genome, Guo *et al*. (2014). Indeed, the binding is strictly regulated by the chromatin context. The simple methylation of the central cytosine of the E box or the methylation of any of the cytosines of the CACGCG sequence can totally abate the binding of the transcription factor to DNA, Perini *et al*. (2005). Thus prediction of the distribution of Myc, along genome of a given cell, must take into account DNA modifications. For what regards the mapping of the Myc factor on repressed/silenced genes, the attempt is even more complicated, since at the moment we have no clue about the existence of DNA consensus sequences that can be used as marks for such an activity. The accurate prediction of the genome distribution of Myc on a given cell genome is of paramount importance, considering that Myc is one of the most powerful oncogenes directly involved in the arising and progression of more than 80% of different types of cancer. Myc regulates about two thousand genes which can be, however, in part different, depending on the type of tissues and tumours. Thus, it is a priority to devise computational tools that may help determine the function of Myc in a specific tumour, possibly in that of single patients, in order to administer personalized and more effective therapeutic treatments. Although several standard bioinformatic tools have been used to address that problem, results have not been particularly effective especially for what regards gene repressed by MYC.

**Fig. 1.**
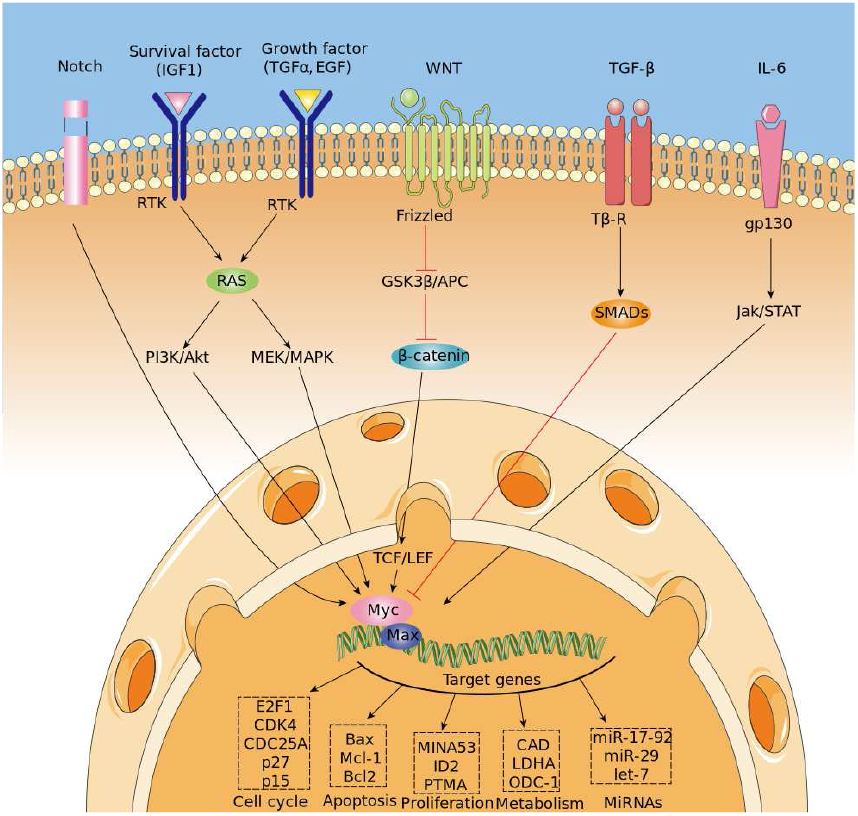
Regulation of MYC expression and binding mechanism

Recently, the scientific community witnessed the outstanding performances of the Deep Learning algorithm in many fields, LeCun *et al*. (2015), Hinton (2018), despite the fact this algorithm was originally developed for image recognition (see Hinton *et al*. (1990), Hinton *et al*. (2006)). The versatilily of the Deep Learning algorithm suggested then new applications in tandem with a specific encoding and representations of DNA sequences. In this work we take a further step and use the DeepBind tool, developed in the Canadian Institute for Advanced Research. With a revolutionary new representation of DNA sequences, going beyond the one-hot-encoding, using data coming via the Chip-Seq databases, researchers were able to predict binding sites and further discern new patterns associated with them, Hassanzadeh and Wang (2016), Trabelsi *et al*. (2019). There are two main advantages in using DeepBind and its refinement DeeperBind. First, DeepBind does not require a large amount of data, in comparison with the other models of DeepRAM and more generally the models coming from Deep Learning architectures. This is a very convenient feature, since large datasets are rarely available, at least not in an homogeneous fashion, in biological databases. Then, the second advantage consists in the fact that DeepBind allows the researchers to view and analyze the filters of the neural network, hence discovering the features that enabled the algorithm to detect the binding sites. For example, ChIPanalyser (see Zabet and Adryan (2015) and refs. therein) and other similar bioconductor packages need consensus sequences and other key information about the transcription factor in order to be able to predict binding sites. On the contrary Deepbind and DeeperBind are able to find old and new consensus sequences associated to a given transcription factor, while at the same time enhancing the performance of binding site recognition. The paves the road to a revolutionary application of the Deep Learning algorithm beyond the supervised tasks. As we shall see in our present work, the analysis of the filters is able to suggest new directions in research by discovering new sequences associated with the MYC-MAX binding dynamics that could be then better explored with further experimental analysis.

## 2 Materials and methods

The DeepBind architecture is a convolutional neural network (CNN) with one convolutional layer followed by a non-linear thresholding, a maxpooling layer and one/two fully connected layers to estimate the intensity of inputs. The DeeperBind model, Hassanzadeh and Wang (2016), is a further development of DeepBind with twice the depth two LSTM (Long short-term memory) recurrent layers stacked on top of the convolutional layers. While the convolutional layers extract the features by applying several PWM-like filters, the RNN (recurrent neural network) captures the sequential dependencies of the sub-motifes identified by the convolutional part of the network.

We performed the training via the DeepRAM tool on the Colab platform. It performs an automatic calibration by training 40 different models whose architecture is summarized in Fig. 2, with randomly sampled hyperparameters and choosing according to the performance of the corresponding model. After the best hyperparameter set is identified, six different models are trained with it. Among this six models DeepRAM will select and save the one with the highest precision. First, we explored the model functionality and behaviour on a relatively small dataset consisting of the collection of 300 sequences from the experiments belonging to just the H1 cell line. Once done with this first exploratory run and selected the best hyperparameters, we proceeded with the training of the model with the given hyperparameters with 12000 sequences obtained with the dataset JFK_4000 obtained by selecting 4000 sequences from the experiments in Chip-Seq. We take sequences of length 301, three times larger than the ones examined by the DeepRAM models in Deepbind and Deeperbind. We also remark that one feature of the model is the random initialization of the filters parameters: consensus sequences are *learned* as the algorithm progresses and determines the weights of our model.

**Fig. 2.**
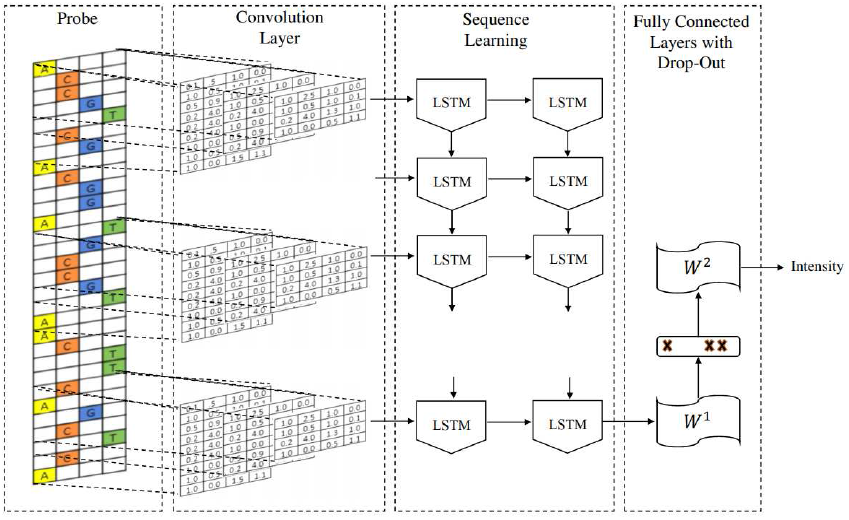
The architecture of the CNN DeeperBind model

We then sorted each of the retrieved experimental data according to the score associated to each peak, that is, the Myc binding site, selecting peaks with the highest score from each experiment. For the purpose of our study we extracted 301nt long sequences from the hg19 human genome assembly. Each sequence was centered on the middle point of the corresponding peak. The sequences thus obtained form the set of positive sequences, in other words, the sequences bound by MYC. Given the set of positive sequences, we generated a set of negative sequences through dinucleotide shuffling, that is the process of randomly shuffling a DNA sequence while preserving the counts of the 16 different dinucleotides. In such a way, the model is prevented from simply relying on the low-level statistics of genomic regions, such as promoters or coding regions, to discriminate positives from negatives.

We summarize in the next two tables the splitting into training, validation and test that we followed for each of our datasets and the composition of our datasets in terms of positive and negative sequences:

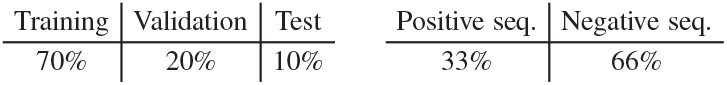

## 3 Results

We run our models with different datasets elucidated in Table 1 and we summarize our results in Table 1 and Fig. 3. Those results have accuracies expressed via AUC (area under the ROC curve), a statistical device, measuring the accuracy of a binary response: the closer to 1 the better is the accuracy. For a thorough explanation of this statistical concept, see Swets (1988) and the expository Fawcett (2006) with refs. therein.

**Table 1.**
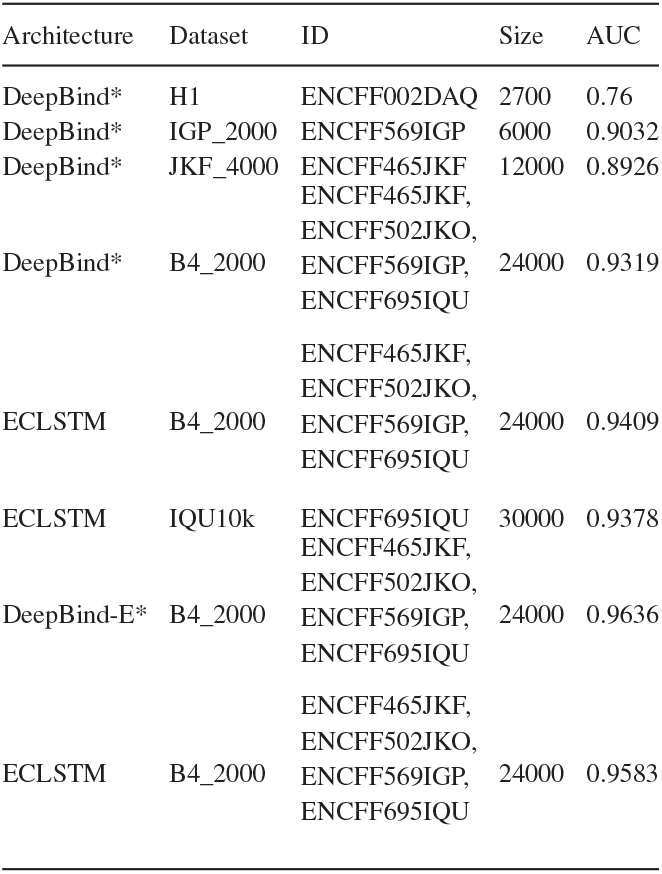
Our DeepBind Models on Myc and AUC

**Fig. 3.**
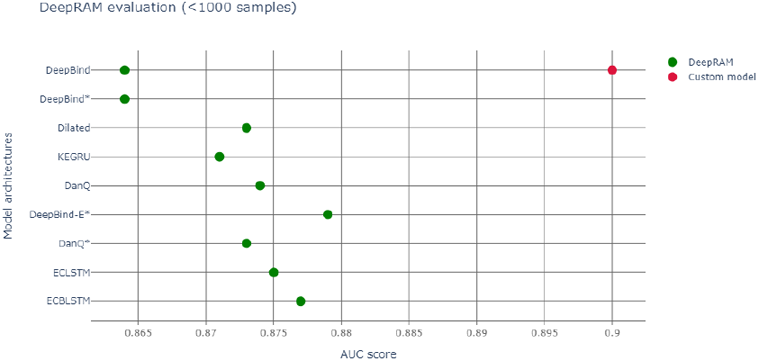
Red dot represents AUC of our model, green dots the various DeepBind architectures (Hassanzadeh and Wang (2016) and Refs. therein)

Table 2 reports the AUC for DeepBind (ref. Hassanzadeh and Wang (2016)), measuring the accuracy in Myc binding site (peak) recognition for the cell line H1-hESC, in parenthesis an estimation of the error.

**Table 2.** AUC of DeepBind on Myc (Hassanzadeh and Wang (2016)) on Training and Test datasets

|  |  |
| --- | --- |
| AUC on training data | 0.9114 (0.0006) |
| AUC on test data | 0.9301 (0.0097) |

So, we notice an accuracy slightly larger than the original Deepbind experiments, though it is not possible to make a full comparison between experiments, due to a different training set size and context. We also report a more appropriate graphic comparison with the other models, with a similar training set size in Fig 3; however those models regarded also other transcription factors besides Myc.

As Fig. 4 shows, at the end of the training, we can examine the learned filters, containing the information the algorithm effectively uses to reach the high accuracy binary classification.

**Fig. 4.**
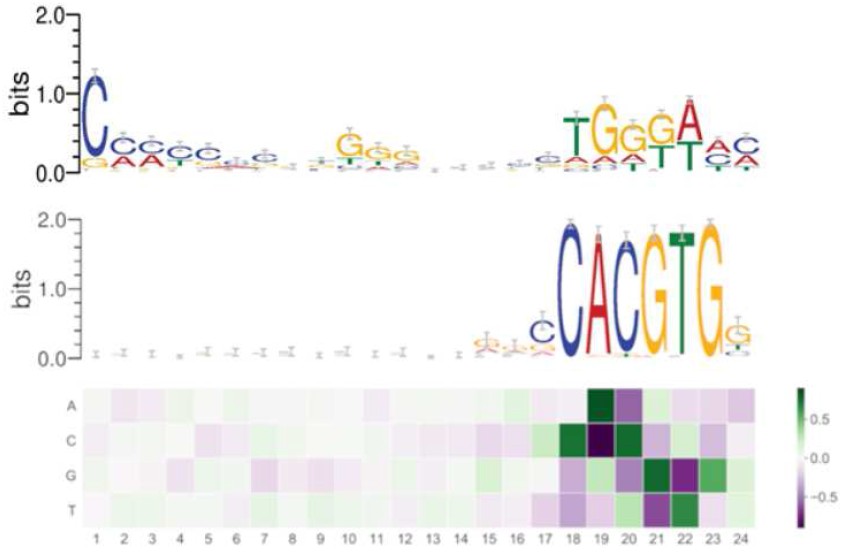
Filter 1 (above), filter 2 (below) expressing the learned consensus sequences in DeeperBind. Coloring for letters marks nucleotides, coloring for the filters indicates the values for the weights, hence their importance (the darker, the higher is the weight).

By analyzing the filters, we discovered two consensus sequences: one for the MYC factor and, surprisingly, another one, that we call “non-canonical”. Such new sequence shares similarities with the one recognized by the RPBJ factor. No other sequence emerges with the same importance as these two, as expressed in Figure 4. Interestingly, the non-canonical consensus sequence can be split into two DNA regions which share some degree of similarity with the DNA consensus sequences respectively recognized by the SP1 transcription factor (C/G rich region) or the MIZ-1 initiation factor (TGGGA), usually used by MYC to exert transcriptional repression. In order to further explore the pattern-learning capabilities of the model, we generated the position weight matrices (PWM) for the two filters in DeeperBind (see Fig. 4), which better captured the consensus sequences. The position weight matrix for DNA analysis is a 4xN matrix commonly used for representation of motifs of length N in biological sequences. They are obtained from sets of aligned sequences in order to check if they are functionally related and they are now an important tool available, through various software implementations, for computational motifs search and discovery. PWM elements are calculated via the position probability matrix (PPM), which counts and normalizes the frequency of each of the four nucleotides for each position in a given sequence, and then checking the distance between their distribution in the given sequence and the distribution in a random sequence with the use of the log likelihood function (related with Kullback Leibler divergence, an effective method to measure the distance between two probability distributions). In our analysis, we first annotated all the sequences we used for our models training, assigning, when possible, a gene to each sequence. Among the 8000 sequences we started with (Table 1), we we were able to identify 5591 genes. We then built two datasets containing the sequences corresponding to the genes whose expression is, respectively, promoted or repressed by Myc. In the end, comparing the annotated sequences with the validated target of Myc transcriptional activation and repression, we obtained 53 sequences (of length 301) corresponding to target transcriptional activation and 21 sequences (of length 301) corresponding to target transcriptional repression. We then tested the two PWM obtained from the models filters on each of the two datasets. We found out that the PWM obtained from the filter (through a sliding procedure) which learned the canonical consensus sequence (Fig. 4, filter 2) had higher affinity (score) for 41 sequences among the 53 transcriptional activation targets compared to the non-canonical one (Fig. 4, filter 1). On the other hand, the non-canonical consensus sequence (Fig. 4, filter 1) had higher affinity for 14 sequences among the 21 transcriptional repression targets as expected. Our comparison is based on assigning a score to the comparison via PWM between the sequence (of length 24) of filter 1 (resp. filter 2) of Fig. 4 and each sequence on each of the 21 (resp. 53) sequences of transcriptional activation (resp. repression) of Myc. In 14 (resp. 41) instances the score of filter 1 is higher than the one obtained with filter 2. We summarize our findings in Table 3.

**Table 3.** Comparison of number of detected consensus sequences in filters 1 and 2.

| Model filters | Consensus seq.<br>(Myc act./repress.)<br>in models filters | No. of transcr. act./<br>repr. seq. in database | No. of transcr. act./ repr.<br>Seq. with higher score for<br>the filter |
| --- | --- | --- | --- |
| Filter 1 | (G/T)GGGCGG<br>(G/A)(G/A)(G/T) | 21 | 14 |
| Filter 2 | CANNTG | 53 | 41 |

These findings go in the direction of the current understanding of Myc binding dynamics and suggest further investigation in both confirming our qualitative findings for the new consensus sequence (Fig. 4, filter 1) and then in trying new models to discover novel consensus sequences both for activation and repression. Our findings support the idea that our in silico approach can help discriminate those DNA sequences that can serve as either Myc binding site for active transcription or sites for indirect association of Myc to genes that are repressed/silenced (Table 3). The analysis, even if performed on a small annotated dataset, highlights the capabilities, not yet fully exploited, of CNN models to extract and generalize the contextual relevance of regulatory sequences. We further speculate that, with more annotated data and the consequently appropriate training, the model could learn more consensus sequences related with the functional role played by a given transcription factor.

## 4 Conclusions

DeeperBind paired with DeepRAM obtaines extraordinary results in predicting binding sites for the MYC factor, reaching an accuracy above 0.96 AUC, when trained with multiple cell lines (see Table 1). We notice a substantial improvement in the accuracy (AUC), when we move from a training set extracted from just one cell line to one coming from multiple (up to six) cell lines: this is possibly linked to the fact that MYC factor is independent from the cell line, hence training with variations enhances the recognizing ability of the algorithm. From the analysis of the filters, we discover that the model does not only learn the consensus sequence CACGTG classically associated to the MYC factor, but learns also other motifs with less significance for the classification task. For example in Fig. 2 we see the consensus sequence TGGGA, associated with the RPBJ factor, suggesting a possible correlation between the presence of MYC and RPBJ factors. We plan to explore this further in the future. In conclusion, we believe that mapping each filter on positive sequences may help to identify which filter is more relevant for the prediction and were it maps on the actual biological sequence. It also can suggest new correlations between different factors as MYC and RPBJ. These capabilities may open the path for novel biological findings without the extensive use of time and resource consuming laboratory essays.

